# Medical relevance of common protein-altering variants in GPCR genes across 337,205 individuals in the UK Biobank

**DOI:** 10.1101/2019.12.13.876250

**Authors:** Christopher DeBoever, AJ Venkatakrishnan, Joseph M Paggi, Franziska M. Heydenreich, Suli-Anne Laurin, Matthieu Masureel, Yosuke Tanigawa, Guhan Venkataraman, Michel Bouvier, Ron O. Dror, Manuel A. Rivas

## Abstract

G protein-coupled receptors (GPCRs) drive an array of critical physiological functions and are an important class of drug targets, though a map of which GPCR genetic variants are associated with phenotypic variation is lacking. We performed a phenome-wide association analysis for 269 common protein-altering variants in 156 GPCRs and 275 phenotypes, including disease outcomes and diverse quantitative measurements, using 337,205 UK Biobank participants and identified 138 associations. We discovered novel associations between GPCR variants and migraine risk, hypothyroidism, and dietary consumption. We also demonstrated experimentally that variants in the β_2_ adrenergic receptor (*ADRB2*) associated with immune cell counts and pulmonary function and variants in the gastric inhibitory polypeptide receptor (*GIPR*) associated with food intake and body size affect downstream signaling pathways. Overall, this study provides a map of genetic associations for GPCR coding variants across a wide variety of phenotypes, which can inform future drug discovery efforts targeting GPCRs.

## Introduction

G-protein-coupled receptors (GPCRs) are allosteric molecular machines that detect extracellular stimuli, such as small molecules, peptides, light, or ions, and in response initiate intracellular signaling pathways. There are over 800 GPCRs in the human proteome that play key roles in diverse physiological functions, and more than one third of FDA-approved medications target this class of receptors^1^. Interest from the pharmaceutical industry and basic science has fueled intensive study of GPCRs, yet there still exist many GPCRs for which the primary physiological function is unknown, and even for well-studied receptors the full range of physiological impacts may not be known. Understanding the roles of GPCRs is particularly relevant because their positioning on the cell surface makes them relatively easy to target with drugs, making it realistic to translate an association into a medication.

The UK Biobank dataset contains genetic and phenotypic data for nearly 500,000 individuals and provides a unique opportunity to study the physiological impact of GPCRs by performing a phenome-wide association study (PheWAS) to identify associations between variants in GPCRs and diverse phenotypes such as immune cell measurements, vital signs, and disease risk^2^. A PheWAS focused on protein-altering variants in GPCRs can connect GPCRs to specific phenotypes and generate novel therapeutic hypotheses as they are likely candidates for causal variants^3^. We can also leverage the increasing number of GPCR protein structures and shared structural scaffold across the GPCR family to identify specific genetic variants that are likely to impact protein structure and explain the genetic associations. Additionally, while the vast array of GPCRs recognize a diverse set of extracellular signals, they share a common set of intracellular binding partners: G proteins and arrestins^4^. G protein and arrestin coupling are directly measurable, providing insight into the molecular impact of GPCR variants.

In this study, we performed a PheWAS for 269 missense and protein-truncating variants in 156 GPCR genes across 275 diverse phenotypes in the UK Biobank, including disease outcomes and quantitative phenotypes, and identified 138 associations at a false discovery rate of 5%. The associations spanned 52 coding variants in 41 GPCRs and 46 quantitative or binary phenotypes, and included both novel and previously reported associations. We identified five associations with binary phenotypes including an association between rs12295710 in *MRGPRE*, a gene whose paralogs are involved in nociception, and migraine^5^. The remaining 133 associations were between coding GPCR variants and quantitative phenotypes including immune cell measurements, body size, pulmonary function, food intake, and others. Five orphan receptors of unknown function had associations with quantitative traits; these orphan GPCRs include *GPR35*, a known drug target for inflammatory bowel disease and cardiovascular disease^6^. We also found associations between variants in GPCR taste receptors known to affect bitter taste perception and self-reported tea and coffee intake. We tested genetic variants in the β_2_ adrenergic receptor (*ADRB2*) associated with several phenotypes for effects on downstream signaling pathways and identified two *ADRB2* haplotypes with differential signaling relative to the most frequent *ADRB2* haplotype. We found associations between rs1800437 (p.Glu354Gln) in the gastric inhibitory polypeptide receptor (*GIPR*) and multiple phenotypes including body size and fresh fruit intake. We assessed the impact of p.Glu354Gln on the major signaling pathways of *GIPR* and observed that they are all blunted for the alternate allele (Gln) relative to the reference allele (Glu). This study links coding variation in GPCRs to a range of diverse phenotypes and identifies novel biology for this important class of drug targets.

## Results

### GPCR genetic associations across 275 phenotypes

To assess the impact of genetic variation in GPCRs on disease risk and other phenotypes, we performed genetic association analyses between GPCR variants and phenotypes for 337,205 individuals of unrelated white British ancestry in the UK Biobank (see Methods for description). Beginning with a list of 226 non-olfactory human GPCRs, we identified 251 missense and 18 protein-truncating variants in 156 human GPCR genes that were genotyped in the UK Biobank arrays, passed quality control filters (Methods), and had minor allele frequency (MAF) greater than 1% (Table S1). We tested for associations between these 269 variants and 146 quantitative phenotypes with at least 3,000 observations and 129 binary phenotypes with at least 2,000 cases in the studied cohort (Methods, Table S1). The quantitative phenotypes include continuous or ordinal phenotypes such as weight, waist size, and forced expiratory volume. The binary phenotypes include the presence or absence of health conditions such as hyperthyroidism, migraine, or lung cancer. Due to power differences between quantitative and binary phenotype association tests, we corrected for multiple hypothesis testing separately for the quantitative and binary phenotypes using the Benjamini-Yekutieli (BY) procedure at FDR rate of 5%. Overall, we identified 52 coding variants in 41 GPCRs that were associated with at least one of 46 quantitative or binary phenotypes for a total of 138 associations (Table S2).

Taking advantage of GPCRs sharing a common fold, we annotated the tested variants with across-family conservation scores, GPCR functional regions, and PolyPhen-2 pathogenicity scores (Methods)^4,7,8^. We compared the variants with and without significant associations to 86,601 rare protein-altering GPCR variants from gnomAD^9^ (MAF<1%) and 318 GPCR variants reported in ClinVar^10^ and found that both the significant and non-significant UK Biobank variants had significantly lower PolyPhen-2 scores than the ClinVar and rare gnomAD GPCR variants (Wilcoxon, p<1×10^−5^), consistent with the fact that the GPCR variants tested here are common (MAF > 1%) variants (Figure S1a-d, Table S3). We did not find a significant difference between the UK Biobank GPCR variants with or without significant associations (Wilcoxon, p=0.69). We also compared family conservation scores for variants in class A GPCRs and found that the conservation scores generally agreed between the different variant sets except that ClinVar variants had higher conservation scores than gnomAD rare variants (Wilcoxon, p=3.8×10^−6^) and that variants without significant associations had lower conservation scores than ClinVar variants (Wilcoxon, p=0.013, Figure S1e, Table S3). Among the 52 variants with significant associations, we identified eight variants with associations that have PolyPhen-2 scores greater than 0.9, four variants located in the ligand binding pocket, and two variants located in the intracellular coupling interface (Table 1). For instance, we identified an association between rs3732378 (p.Thr280Met, MAF=17.3%) in *CX3CR1* and hypothyroidism (p=1.3×10^−7^, OR=1.07, 95% CI: 1.05-1.10). rs3732378 has a PolyPhen-2 score of 0.774 and is located in the binding pocket of CX3CR1. This GPCR is the sole receptor for the ligand fractalkine (CX3CL1), a chemokine with anti-apoptotic properties that has been implicated in oncogenesis^11,12^. We also identified associations between rs3732378 and monocyte counts, monocyte percentages, and lymphocyte counts (Table S2), consistent with previous studies^13^. Expression of CX3CR1 is upregulated in the absence of triiodothyronine (T3) by thyroid receptors TRα1 and TRβ1 with mutations observed in human hepatocellular carcinoma suggesting that thyroid receptors may regulate CX3CR1 in some circumstances ^14^. We also identified associations between rs2229616 (p.Val103Ile, MAF=2.0%) and body size phenotypes and immature reticulocyte fraction. rs2229616 was recently reported as a gain-of-function variant that exhibits significant bias toward β-arrestin signaling^15^. Overall, these results indicate that although common variants are generally not expected to have a high impact on proteins due to evolutionary constraint, some common coding GPCR variants are associated with phenotypic variation and likely impact on protein function (Table 1).

**Table 1.** GPCR variants with significant associations and PolyPhen-2 scores greater than 0.9 or known functional annotations. “Number of Associations” indicates the number of significant associations identified for each variant in this study across both quantitative and binary phenotypes.

| Gene | Variant | MAF | HGVSp | PolyPhen-2 | Function | Number of Associations |
| --- | --- | --- | --- | --- | --- | --- |
| *GIPR* | rs1800437 | 19.4% | p.Glu354Gln | 1 |  | 13 |
| *ADRB2* | rs1800888 | 1.5% | p.Thr164Ile | 0 | Binding | 6 |
| *ACKR2* | rs2228467 | 6.1% | p.Val41Ala | 0.958 |  | 5 |
| *MC4R* | rs2229616 | 2.0% | p.Val103Ile | 0.025 | Binding | 5 |
| *CX3CR1* | rs3732378 | 17.3% | p.Thr280Met | 0.774 | Binding | 4 |
| *P2RY2* | Affx-5881601 | 24.0% | p.Arg312Ser | 0.015 | Coupling | 4 |
| *CELSR3* | rs3821875 | 11.2% | p.Ser805Thr | 0.999 |  | 4 |
| *LGR4* | rs34804482 | 2.6% | p.Asp844Gly | 1 |  | 2 |
| *P2RY13* | rs1466684 | 18.0% | p.Thr179Met | 0.001 | Binding | 2 |
| *GPR35* | rs3749171 | 18.2% | p.Thr139Met | 0.956 |  | 2 |
| *GPR45* | rs35946826 | 14.3% | p.Leu312Phe | 1 |  | 1 |
| *NPFFR1* | rs3812694 | 5.9% | p.Ile145Leu | 0.999 | Coupling | 1 |
| *S1PR3* | rs34075341 | 3.6% | p.Arg243Gln | 1 |  | 1 |

### Associations with binary medical phenotypes

We identified five associations between GPCR variants and binary phenotypes (BY-adjusted p < 0.05, Table 2, Table S2). We identified a novel association between rs12295710 (p.Gly15Ser, MAF=47.1%) in *MRGPRE*, a member of the Mas-related receptor family, and migraine (p=3.8×10^−8^, OR=1.07, 95% CI: 1.05-1.10). Neither this variant nor other variants in linkage disequilibrium (LD) with it (R^2^>0.8, 1000 Genomes British in England and Scotland (GBR)) have been previously reported as associated with migraine in the GWAS Catalog, in a meta-analysis of migraine GWAS including 375,000 individuals, or in a study of broadly-defined headaches in the UK Biobank^16–18^. Though the endogenous ligand of MRGPRE is unknown, Mas-related receptors are expressed in nociceptive sensory neurons and play a role in pain response. MRGPRE is expressed throughout the brains of macaques, mice, and humans including sensory neurons in mice^5,19,20^. Suggestive associations between variants in *MRGPRE* and white matter mean diffusivity and brain lesion distribution in multiple sclerosis have been reported indicating a possible role for *MRGPRE* in neurological disorders^21,22^.

**Table 2.**
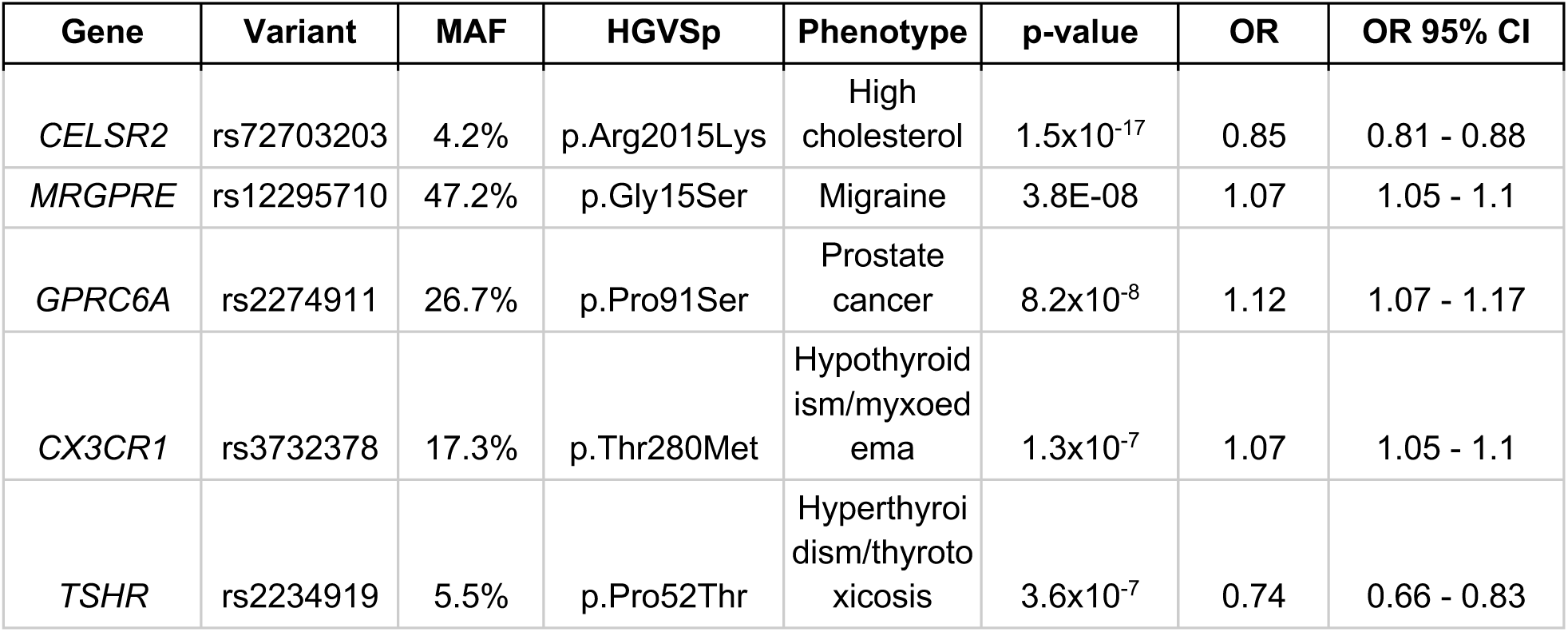
Significant associations between coding GPCR variants and disease outcomes.

We found an association between rs2234919 (p.Pro52Thr, MAF=5.5%) in the Thyroid Stimulating Hormone Receptor (*TSHR*) and hyperthyroidism (p=3.6×10^−7^, OR=0.74, 95% CI: 0.66-0.83). Previous studies have found conflicting evidence regarding the association between rs2234919 and hyperthyroidism^23^, and the variant is flagged as likely benign in ClinVar consistent with our finding that the minor allele lowers hyperthyroidism risk. We also identified an association between rs72703203 (p.Arg2015Lys, MAF=4.2%) in *CELSR2* and high cholesterol (p=1.5×10^−17^, OR=0.85, 95% CI: 0.81-0.88). Variants in or near *CELSR2* have been previously associated with cholesterol, response to statins, lipoprotein-associated phospholipase A2 activity, and coronary artery disease^24–28^. While rs72703203 is not in high LD with previously reported variants (Figure S2), it is not significant (p=0.21) in a model that includes rs12740374, a 3’ UTR variant in *SORT1* proposed as the causal variant in this region^29^. We identified an association between rs2274911 (p.Pro91Ser, MAF=26.7%) in *GPRC6A* and prostate cancer (p=8.2×10^−8^, OR=1.12, 95% CI: 1.07-1.17). rs2274911 is in LD with rs339331 (R^2^=0.846, GBR) which has previously been associated with prostate cancer, and rs2274911 is not significant in a model that includes rs339331 as a covariate (p=0.47)^30–32^.

### Associations with quantitative phenotypes

We identified 133 quantitative phenotype associations for 49 protein-altering variants in 38 genes and 41 distinct phenotypes (BY-adjusted p < 0.05, Figure 1A-B, Table S2). 48 of these associations are present in this GWAS Catalog or in LD (R^2^>0.8, GBR) with GWAS Catalog associations^16^. We identified novel associations for variants in three GPCRs with no entries in the GWAS catalog: rs7570797 in *PROKR1* (p.Ser40Gly, MAF=4.6%) was associated with plateletcrit (p=1.59×10^−7^, *β*=-0.023, 95% CI: −0.032 - −0.015); rs1466684 in *P2RY13* (p.Thr179Met, MAF=18.0%) was associated with neutrophil count (p=1.68×10^−6^, *β*=0.016, 95% CI: 0.0094-0.023); and rs4274188 in *MRGPRX3* (p.Asn169Asp, MAF=23.4%) was associated with standing height (p=7.69×10^−6^, *β*=0.0090, 95% CI: 0.0051-0.013). We stratified the significant quantitative phenotype associations into the following phenotype categories (where n indicates the number of phenotypes with at least one association for each category): immune cell measurements (n=17), body size (n=6), lung function (n=6), food intake (n=4), physical ability (n=3), urine biomarkers (n=2), vital signs (n=2), and intelligence (n=1) (Table S1). We describe these associations in the sections below.

**Figure 1.**
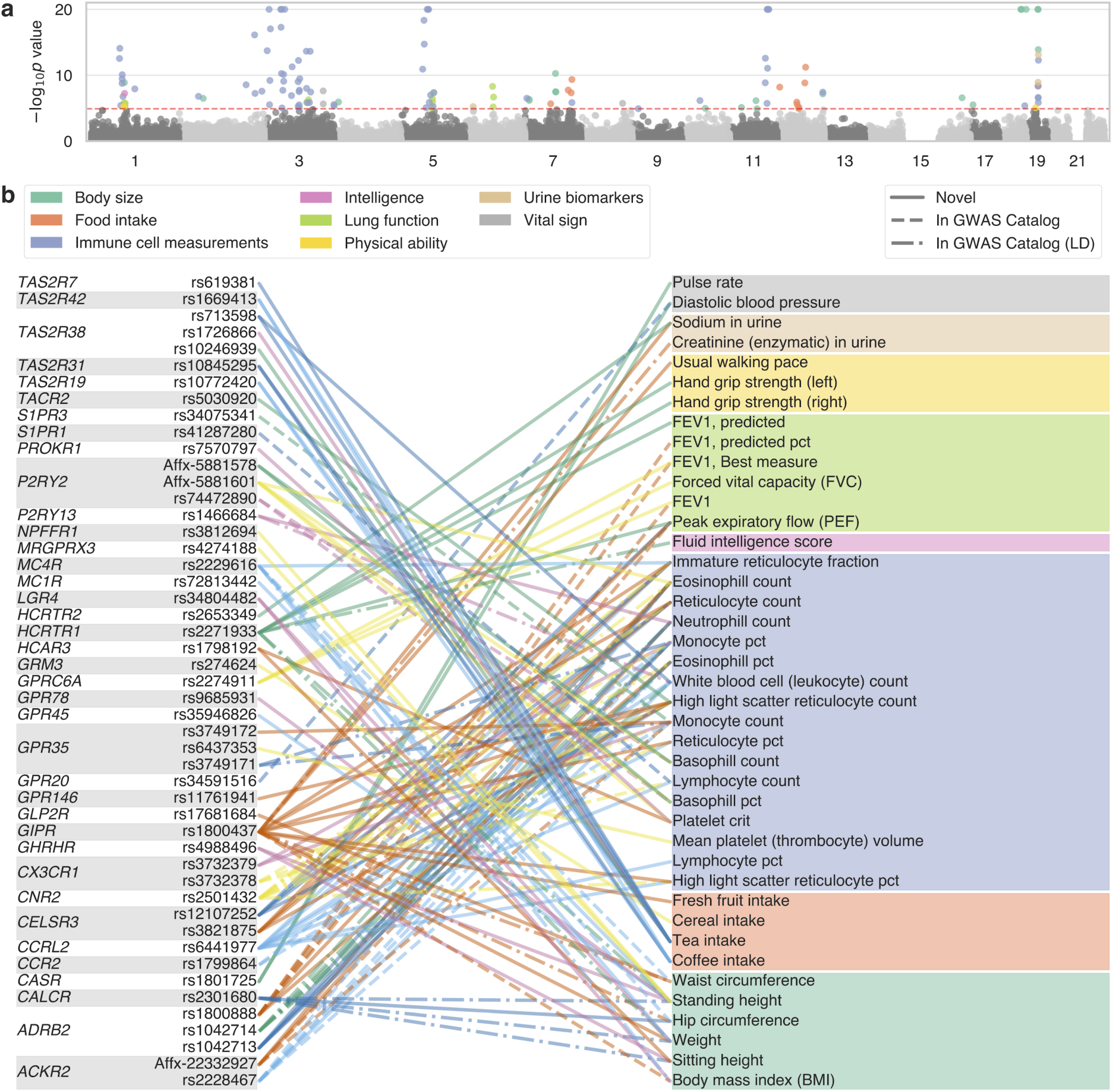
(a) Manhattan plot for quantitative phenotype associations across 269 variants and 146 phenotypes. Dashed red line indicates significance threshold after applying Benjamini-Yekutieli procedure. Scatter point colors indicate phenotype category (see legend in (b)) for significant associations. P-values less than 1×10^−20^ are plotted at 1×10^−20^. (b) Line plot showing associations between coding variants in indicated GPCRs (left) and quantitative phenotypes (right). Colors indicate phenotype category. Dashed lines indicate associations that are present in the GWAS Catalog and dot-dashed lines indicate that the variant is in LD with a variant associated with the phenotype in the GWAS Catalog.

### Genetic variation in orphan GPCRs associated with quantitative phenotypes

The 269 variants tested for associations included 44 variants in 26 orphan GPCRs whose ligands and function are generally unknown^33^. We identified quantitative phenotype associations for variants in five orphan receptors (Table 3). We found associations between rs3749171 in *GPR35* (p.Thr139Met, MAF=18.2%) and monocyte count (p=7.5×10^−17^, *β*=-0.024, 95% CI: - 0.031 - −0.019) and monocyte percentage (p=5.7×10^−8^, *β*=-0.25, 95% CI: −0.016 - −0.034). rs3749171 is in LD with the 5’ UTR variant rs34236350 (R^2^=1, GBR) that has previously been reported as associated with monocyte count (Figure S3) ^13^. rs3749171 is in high LD with the intron variant rs4676410 (R^2^=0.82, GBR), and these two variants have previously been associated with ankylosing spondylitis, ulcerative colitis/inflammatory bowel disease, and pediatric autoimmune diseases (Figure S3)^34–40^. We also found associations between rs6437353 (p.Arg13His, MAF=45.6%) and mean platelet (thrombocyte) volume (p=2.8×10^−9^, *β*=-0.021, 95% CI: −0.028 - −0.014) and between rs3749172 (p.Ser294Arg, MAF=42.6%) and monocyte count (p=2.6×10^−8^, *β*=0.013, 95% CI: 0.0084-0.017). These variants are not in LD with rs3749171 or rs34236350 (Figure S3), and neither these variants nor variants in LD with them have been previously reported as associated with these phenotypes in the GWAS Catalog^16^.

**Table 3.** Significant quantitative phenotype associations for orphan GPCRs.

| Gene | Variant | MAF | HGVSp | Phenotype | p-value | BETA | BETA 95% CI |
| --- | --- | --- | --- | --- | --- | --- | --- |
| *GPR35* | rs3749171 | 18.2% | p.Thr139Met | Monocyte count | $7.5 \times 10^{-17}$ | -0.025 | -0.031 - -0.019 |
| *GPR35* | rs6437353 | 45.6% | p.Arg13His | Mean platelet (thrombocyte) volume | $2.8 \times 10^{-9}$ | -0.021 | -0.028 - -0.014 |
| *GPR35* | rs3749172 | 42.6% | p.Ser294Arg | Monocyte count | 2.6x10<sup>-8</sup> | 0.013 | 0.0084 - 0.017 |
| *GPR35* | rs3749171 | 18.2% | p.Thr139Met | Monocyte percentage | 5.7x10<sup>-8</sup> | -0.025 | -0.034 - -0.016 |
| *GPR146* | rs11761941 | 14.8% | p.Gly11Glu | High light scatter reticulocyte count | 3.1x10<sup>-7</sup> | -0.017 | -0.023 - -0.01 |
| *GPR45* | rs35946826 | 14.3% | p.Leu312Phe | Standing height | 3.2x10<sup>-7</sup> | -0.012 | -0.017 - -0.0077 |
| *GPR78* | rs9685931 | 11.3% | p.Arg342His | Standing height | 1.1x10<sup>-6</sup> | 0.013 | 0.0078 - 0.018 |
| *GPR20* | rs34591516 | 4.6% | p.Gly313Ser | Diastolic blood pressure, automated reading | 1.8x10<sup>-6</sup> | 0.028 | 0.017 - 0.04 |

We found associations for missense variants in orphan GPCRs *GPR146, GPR20, GPR45*, and *GPR78* as well (Table 3). The association between rs34591516 (p.Gly313Ser, MAF=4.6%) in *GPR20* and diastolic blood pressure (p=1.8×10^−6^, *β*=0.028, 95% CI: 0.017-0.04) agrees with a previous study that found the same variant associated with diastolic and systolic blood pressure^41^. We find that rs9685931 (p.Arg342His, MAF=11.3%) in *GPR78* is associated with standing height (p=1.1×10^−6^, *β*=-0.012, 95% CI: −0.0077 - −0.017). A different variant in the 3’ UTR of *GPR78*, rs3775887, is in weak LD with rs9685931 (LD=0.18, GBR) and has previously been associated with FVC. Overall, these associations indicate potential functions or relevant pathways for these orphan GPCRs.

### Genetic variation in GPCRs associated with food and beverage intake

We identified 12 associations between coding variants in GPCRs and food intake phenotypes. We found that variants in several taste receptors were associated with either coffee or tea intake (Table 4). Variants in the bitter taste receptor genes *TAS2R19, TAS2R31*, and *TAS2R42* were associated with coffee intake, though the variants in *TAS2R19* and *TAS2R31* are in high LD (R^2^=0.9, GBR). rs10772420 (p.Arg299Cys, MAF=46.8%) in *TAS2R19* has been shown to affect perception of bitterness, though it is unclear whether rs10772420, rs10845295 (p.Arg35Trp, MAF=48.6%) in *TAS2R31*, or a different variant is causal^42,43^. We also found associations between variants in the taste receptors *TAS2R19, TAS2R31, TAS2R38, TAS2R42*, and *TAS2R7* and tea intake. The same variants in *TAS2R19* and *TAS2R31* that are in high LD and were associated with coffee intake are also associated with tea intake. The variants in *TAS2R7* and *TAS2R42* associated with tea intake are in moderate LD (R^2^=0.59, GBR). The variants in *TAS2R38* that are associated with tea intake are in high LD and are known to affect the ability to taste phenylthiocarbamide (PTC) and 6-n-propylthiouracil (PROP) (Figure S4)^44,45^. The ability to taste PTC follows a dominant model of inheritance where the alternate “taster” allele C is dominant to the “non-taster” reference T allele for rs10246939^44^. We therefore fit additive and genotypic models for tea intake and the variants in *TAS2R38* to test whether the observed association with tea intake is dominant. Interestingly, we found that the association between variants in *TAS2R38* and tea intake is consistent with an additive effect where each additional copy of the taster C allele is associated with increased tea intake (Figure 2A). We did find evidence, however, that the association between rs10772420 in *TAS2R19* and coffee intake is non-additive, where only one copy of the alternate allele accounts for most of the decrease in coffee intake (Figure 2B). We found weaker evidence that the association between variants in *TAS2R19* and tea intake is non-additive, where one copy of the alternate alleles accounts for most of the increase in tea intake (Figure 2C). As noted above, rs10772420 is in LD with the rs10845295 in *TAS2R31* (0.895, GBR). Overall, we identified three independent associations between variants in taste receptors and tea intake and two independent associations for coffee intake.

**Table 4.**
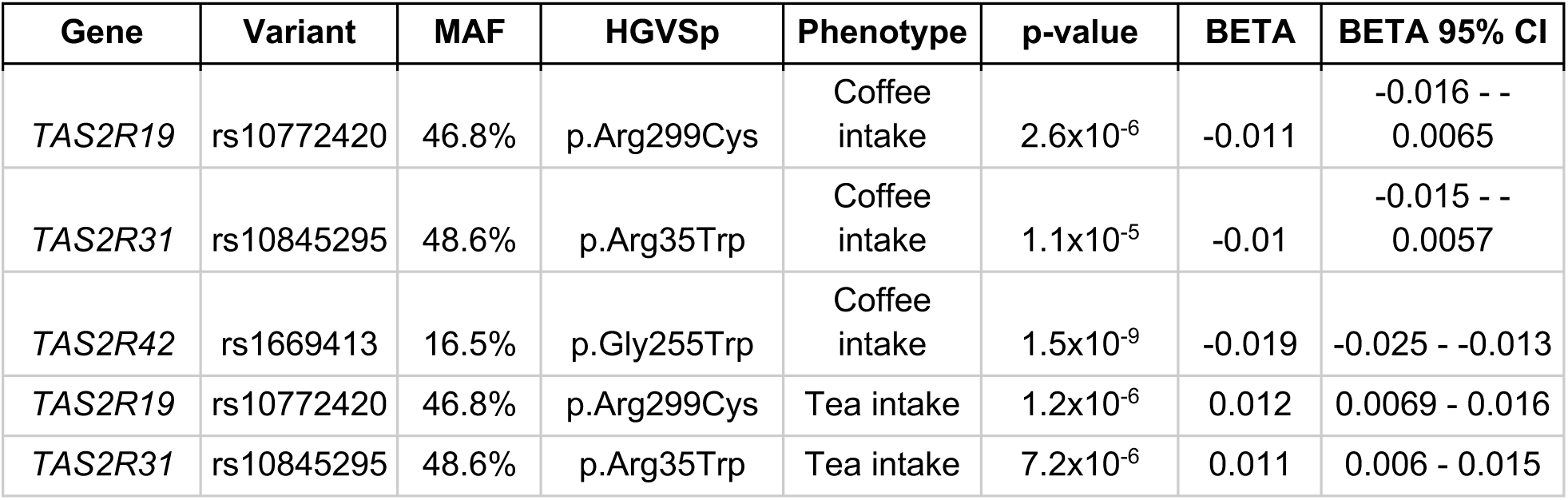
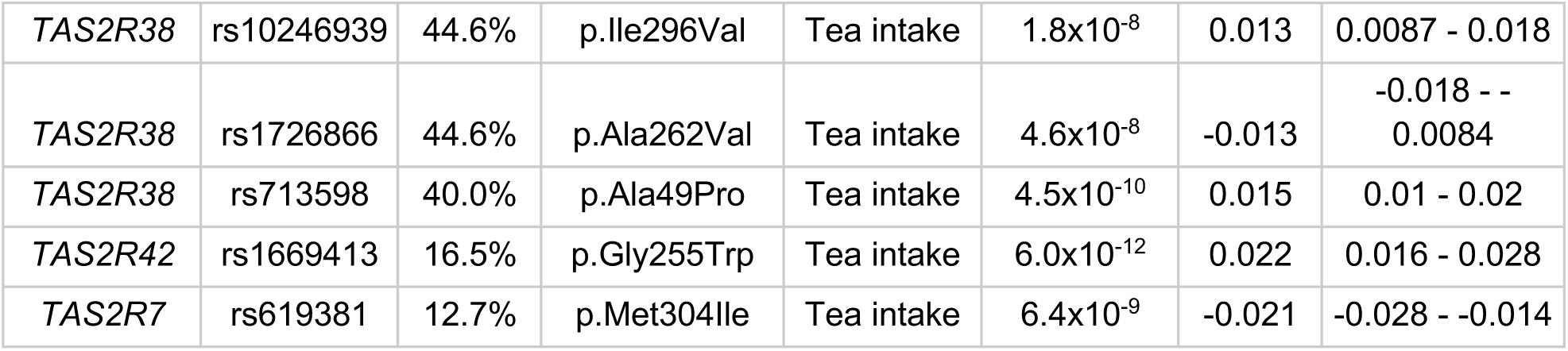
Associations between variants in taste receptors and coffee or tea intake.

**Figure 2.**
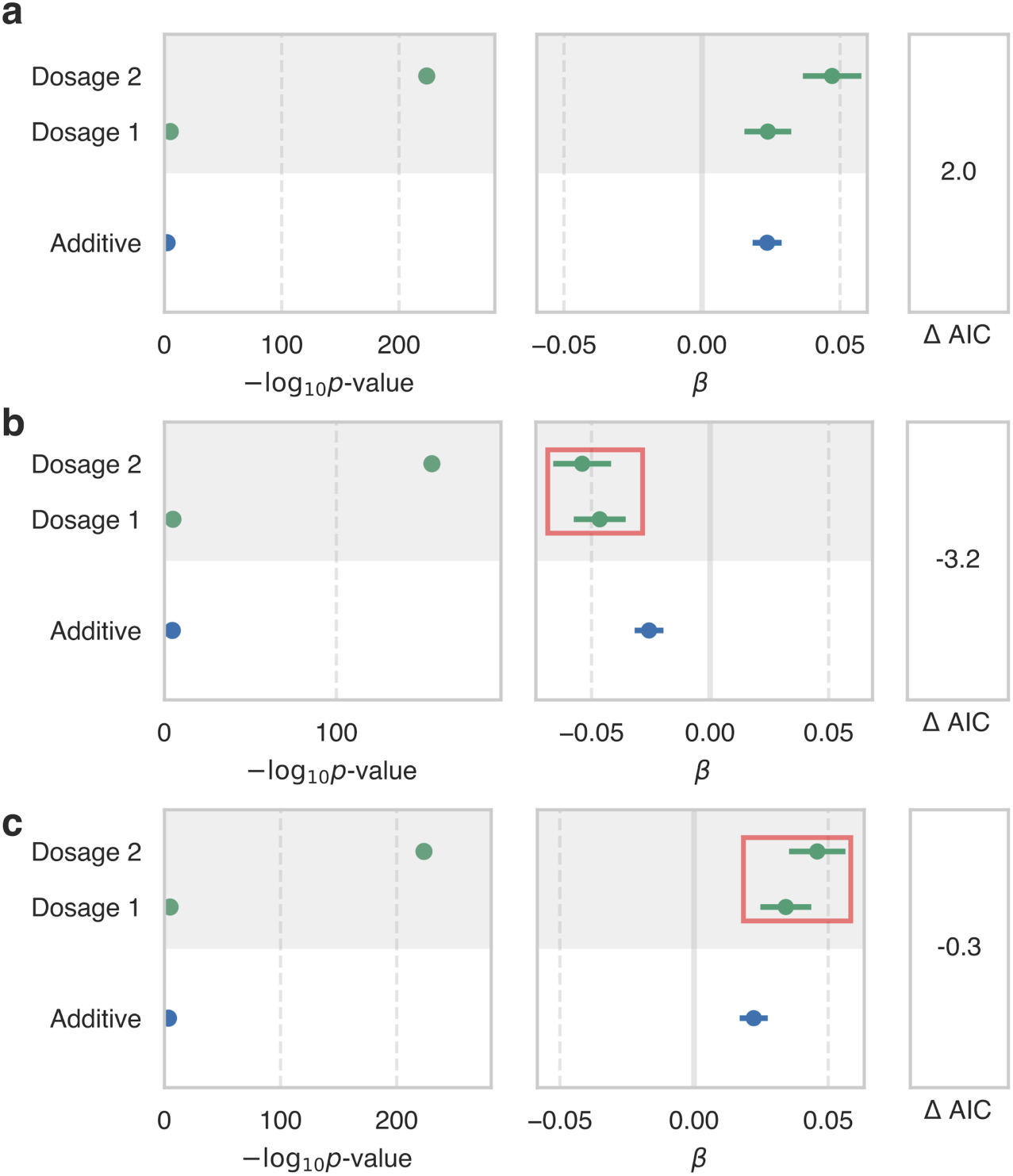
-log_10_p-values and effect sizes for additive (white background) and genotypic (grey background) models of association between (a) rs10246939 in *TAS2R38* and tea intake, (b) rs10772420 in *TAS2R19* and coffee intake, and (c) rs10772420 in *TAS2R19* and tea intake. The genotypic model includes separate terms for heterozygous and homozygous alternate genotypes. Red boxes indicate non-additive effects.ΔAIC is the difference in AIC between the two models (genotypic minus additive).

We also observed two associations between non-taste-related GPCRs and self-reported food intake. We found that rs1800437 (p.Glu354Gln, MAF=19.4%) in *GIPR* is associated with fresh fruit intake (p=4.2×10^−9^, *β*=0.017, 95% CI: 0.011-0.023). *GIPR* is a receptor of gastric inhibitory polypeptide (GIP), which stimulates insulin release in the presence of elevated glucose^46^. Mice lacking *GIPR* exhibit higher blood glucose levels and impaired initial insulin response following oral glucose load^47^. We also identified several other associations for rs1800437 in *GIPR* (Figure 1b, Table S2) that agree with reported associations in the GWAS Catalog including associations with body mass index, weight, waist circumference, and hip circumference, though the reported association with fresh fruit intake has not been previously reported^16^. To explore the impact of rs1800437 (p.Glu354Gln) on the GIP receptor signaling profile, we used bioluminescence resonance energy (BRET)-based assays to monitor β-arrestin2 recruitment and G protein activation^48,49^. It has recently been shown that *GIPR* can promote the activation of Gs, G15 and all Gi/o family members^50^. Dose-response curves for the alternate Gln allele in response to endogenous ligand (GIP) show that maximal activation (Emax) compared to the reference Glu allele was significantly decreased for all Gα pathways tested (65 ± 5 % for Gs, 36 ± 27 % for Gi2, 45 ± 17% for GoB, 55 ± 6% for Gz and 23 ± 6 % for G15) and that β-arrestin recruitment is reduced (statistical test was not performed because parameters could not be extrapolated from unsaturated curve) (Figure S5a,c). The receptor carrying the alternate Gln allele was expressed at a similar or higher cell surface density compared to the receptor with the reference Glu allele as confirmed by enzyme-linked immunosorbent assay (ELISA, Figure S5b). The blunted signaling response for the rs1800437 alternate allele may contribute to the association of rs1800437 with food intake and body size phenotypes.

We also found that a splice donor variant rs274624 (MAF=37.0%) in *GRM3* is associated with cereal intake (p=2.1×10^−6^, *β*=-0.011, 95% CI: −0.015 - −0.0064). *GRM3* encodes a metabotropic glutamate receptor that is involved in brain function. Missense variants in *GRM3* have been previously associated with cognitive performance and self-reported math ability, and non-coding variants near *GRM3* have been associated with several cognitive and psychiatric phenotypes including schizophrenia and neuroticism^16,51^. A non-coding variant near *GRM3* has also been associated with body mass index, but no associations between variants in or near *GRM3* and food intake have been reported previously^52^. These results demonstrate that genetic variation in GPCRs can be linked to food intake using self-reported intake measurements in a population biobank.

### Genetic variation in *ADRB2* alters arrestin signaling

We identified associations between three missense variants in *ADRB2* and immune cell counts and percentages as well as forced expiratory volume in 1-second (FEV1) and peak expiratory flow (PEF) (Table 5). Several of these associations have been reported previously in the GWAS Catalog or are in LD with reported associations, though associations between rs1800888 (p.Thr164Ile, MAF=1.5%) and eosinophil counts (p=5.6×10^−6^, *β*=0.045, 95% CI: 0.026-0.065) and eosinophil percentage (p=4.3×10^−8^, *β*=0.061, 95% CI: 0.039-0.082) were not in the GWAS Catalog. Similarly, the associations between rs1042713 (p.Gly16Arg, MAF=36.1%) and immune cell counts and percentages were not in the GWAS Catalog or in LD with associations in the GWAS Catalog (Table 5)^16^. Since rs1042713 and rs1042714 are in moderate LD (R^2^=0.469, GBR), we fit models for eosinophil counts, eosinophil percentage, monocyte percentage, and neutrophil count using all three variants in *ADRB2*. For all four phenotypes, only one of rs1042713 and rs1042714 were significant (p<0.05) indicating that these two variants likely do not have independent effects on these immune cell phenotypes. rs1800888 remained significant (p<0.05) conditioned on both rs1042713 and rs1042714 for eosinophil counts and percentages.

**Table 5.**
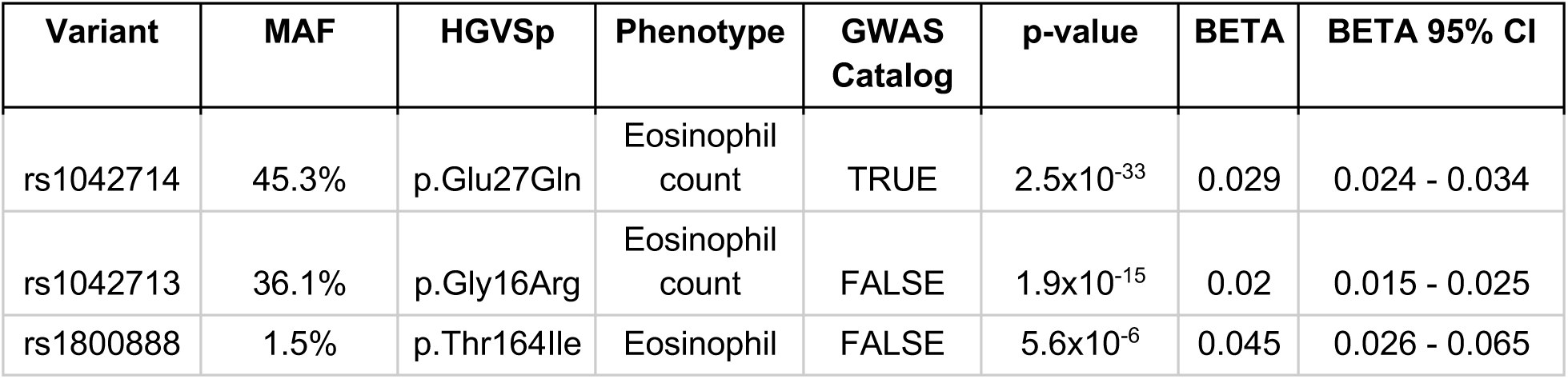
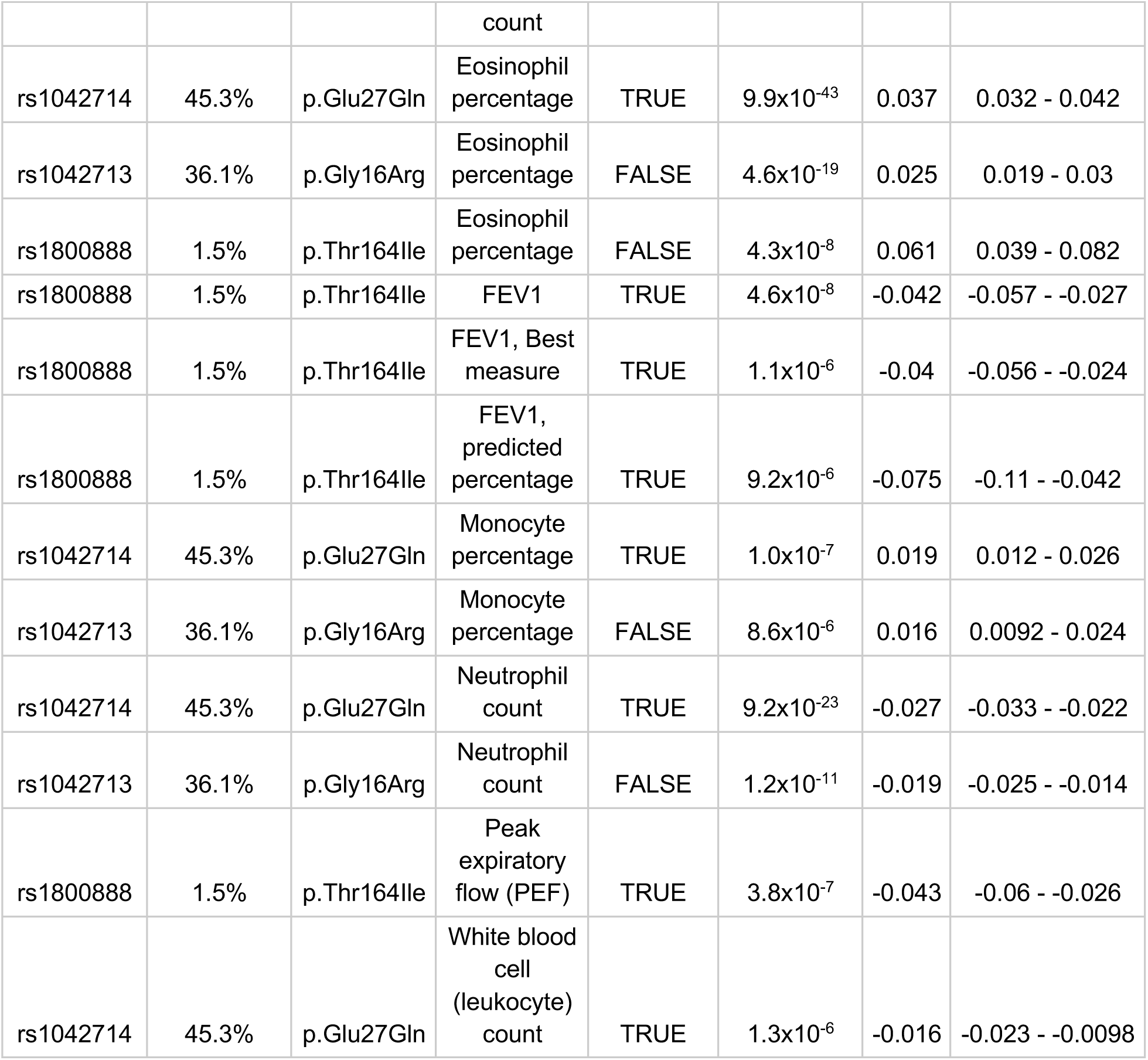
Associations between missense variants in *ADRB2* and quantitative phenotypes. FEV1 is forced expiratory volume in one second.

To explore how the genetic variants in *ADRB2* associated with phenotypes in the UK Biobank impact the function of the receptor, we assayed cAMP signaling, which is downstream of Gs, and *β*-arrestin-1 and *β*-arrestin-2 recruitment, upon stimulation with the endogenous ligands epinephrine (epi) and norepinephrine (norepi) for the four common *ADRB2* haplotypes for rs1042713, rs1042714, and rs1800888: RQT, GET, GQI, and GQT. We refer to the *ADRB2* haplotypes by the residues at the three variable positions; for instance, the most common haplotype among the 1000 Genomes GBR population is RQT (GBR frequency=41.8%) in which rs1042713 codes for arginine (Arg16Gly), rs1042714 codes for glutamine (Gln27Glu), and rs1800888 codes for threonine (Thr164Ile). We found that two of the haplotypes displayed significantly different signaling profiles than the most frequent haplotype RQT. The GET haplotype (GBR frequency=39.6%) retained full arrestin recruitment but significantly affected Gs signaling; for epi, GET has a decreased pEC50 in the cAMP assay (p=0.02, Tukey’s test), indicating that a higher concentration of ligand is required to induce the same level of signaling through cAMP (Figure S6, Table S4). Meanwhile, the GQI haplotype (GBR frequency=2.2%) displayed decreased arrestin pEC50 and full Gs signaling relative to RQT (p < 0.004, Tukey’s test). These differences in signaling between *ADRB2* haplotypes may be responsible for the observed associations between *ADRB2* variants and immune cell counts and pulmonary function.

## Discussion

This study provides a systematic catalog of associations between coding variants in GPCRs and 275 diverse phenotypes in the UK Biobank. We replicated known associations between variants associated with disease risk and quantitative phenotypes and identified novel associations that in some cases indicate novel functions for GPCRs. The association between rs12295710 in *MRGPRE* and migraine is particularly interesting given that the genes in this family play a role in sensing pain^5^. We found that rs3749171 in *GPR35* is associated with monocyte counts and percentage and is in LD with *GPR35* variants associated with ankylosing spondylitis, inflammatory bowel disease, and pediatric autoimmune diseases^34–40^. Interestingly, we also found a novel association between rs3749172 in *GPR35* and monocyte counts, though this variant does not appear to be in LD with loci previously associated with diseases. The novel associations between GPCR taste receptors and self-reported tea and coffee intake demonstrate the utility of questionnaire data in population biobanks, even for relatively difficult-to-estimate phenotypes like food intake.

We tested signaling for *ADRB2* haplotypes for three genetic variants that were associated with immune counts and pulmonary function and found that the GET *ADRB2* haplotype had decreased Gs signaling (as measured by cAMP levels) relative to the RQT haplotype and that the GQI haplotype had decreased arrestin recruitment relative to the RQT haplotype. The observed genetic association between decreased forced expiratory volume and the rs1800888 allele that codes for isoleucine and our results showing that GQI has decreased arrestin signaling might inform the role of *ADRB2* in pulmonary function. Since arrestin recruitment induces *ADRB2* desensitization, and, somewhat paradoxically, *ADRB2* agonist activity is required for the development of asthma in mouse models, it is possible that the GQI haplotype may decrease lung function by reducing agonist-induced *ADRB2* desensitization^53,54^. It is also possible, however, that the acute differences observed in these experiments may differ from long-term effects of differential signaling associated with genetic variants.

Using BRET-based assays that monitor G protein activation or β-arrestin2 recruitment, we showed that the rs1800437 (p.Glu354Gln) *GIPR* alternate allele has blunted signaling for all the pathways tested (Gs, Gi2, GoB, G15 and β-arr2) relative to the reference Glu allele. These results indicate that the alternate Gln allele leads to a general signaling dysfunction that is not specific to a particular pathway and are consistent with a previous study reporting a decreased cAMP signaling^55^. However, some studies have reported either similar responses^56,57^ or even enhanced responses^58,59^ for cAMP accumulation or β-arrestin2 recruitment^59^. It should nonetheless be noted that, in contrast with our study, cell-surface expression of both the reference receptor and the p.Glu354Gln variant receptor was not monitored for each functional assay in many of these reports, complicating the interpretation. Binding affinity was always reported to be identical between the reference Glu allele and the p.Glu354Gln variant^58,59^, suggesting that the diminished signaling of the p.Glu354Gln variant is not a consequence of a lower binding affinity. Moreover, in our study, diminution in signaling efficacy is not explained by reduced cell-surface expression since the receptor carrying the alternate Gln allele was expressed at similar or higher levels than the receptor carrying the reference Glu allele. It is likely that the phenotypes associated with the p.Glu354Gln *GIPR* variant result from a general reduction in signaling rather than from biased signaling, *i.e.* an alteration in the signaling profile favoring or disadvantaging one signaling pathway over another^60^.

The genetic associations reported here link GPCRs to phenotypes and will be useful for assessing the function of GPCRs and potential effects of modulating these genes with therapeutics. Future studies utilizing exome or genome sequencing that can better ascertain rare variants in these genes will likely identify new associations that are not observed for common variants that have faced stronger selective pressure. In particular, studies that focus on populations with unique genetic histories such as founder populations or groups with high consanguinity offer an important opportunity to identify high-impact, rare variants in GPCRs that may have large impacts on phenotypic variation and disease risk.

## Methods

### Quality Control of Genotype Data

We used genotype data from UK Biobank dataset release version 2 for all aspects of the study. To minimize the impact of cofounders and unreliable observations, we used a subset of individuals that satisfied all of the following criteria: (1) self-reported white British ancestry, (2) used to compute principal components, (3) not marked as outliers for heterozygosity and missing rates, (4) do not show putative sex chromosome aneuploidy, and (5) have at most 10 putative third-degree relatives. These criteria are reported by the UK Biobank in the file “ukb_sqc_v2.txt” in the following columns respectively: (1) “in_white_British_ancestry_subset,” (2) “used_in_pca_calculation,” (3) “het_missing_outliers,” (4) “putative_sex_chromosome_aneuploidy”, and (5) “excess_relatives.” We removed 151,169 individuals that did not meet these criteria. For the remaining 337,205 individuals, we used PLINK v1.90b4.4 76 to compute the following statistics for each variant: (a) genotyping missingness rate, (b) p-values of Hardy-Weinberg test, and (c) allele frequencies (calculated separately for the two genotyping arrays used by the UK Biobank).

### GPCR Variants

#### UK Biobank Variant Annotation

Variants on the UK Biobank arrays were annotated using VEP version 87 and the GRCh37 reference^61^. Protein truncating variants were predicted using the VEP LOFTEE plugin (https://github.com/konradjk/loftee). Linkage disequilibrium (LD) was calculated for the GBR British population from 1000 Genomes Phase 3 (Version 5) using LDmatrix from LDlink (https://ldlink.nci.nih.gov/) unless otherwise stated^62^.

#### GPCR Gene Definition

A list of GPCRs was obtained from the International Union of Basic and Clinical Pharmacology (IUPHAR) website in December 2017 (Table S1). This list excludes olfactory GPCRs.

#### GPCR Variant Filtering

We identified 1,263 coding variants in GPCRs that (1) were genotyped on the UK Biobank genotyping arrays, (2) not filtered out during quality control of the genotyping data, (3) were not in the major histocompatibility complex (chr6:28477797-33448354)^63^, and (4) were annotated as one of the following variant types by VEP: missense_variant, frameshift_variant, stop_gained, splice_region_variant, splice_donor_variant, splice_acceptor_variant, stop_lost, inframe_deletion, start_lost, inframe_insertion, incomplete_terminal_codon_variant. We filtered the variants to include only those with minor allele frequency greater than 1% in the 337,205 white British subjects used in this study resulting in 269 variants in 156 genes (Table S1). 251 of the 269 variants are missense variants and the remaining 18 variants are predicted protein truncating variants.

#### GPCR Variant Annotation

86,601 rare protein-altering GPCR variants were obtained from from gnomAD (MAF<1%) and 318 GPCR variants were obtained from ClinVar^10,64^. Family conservation scores were created using multiple sequence alignments from GPCRdb^7^. The scores were defined as the frequency of the most common residue at each structurally equivalent position. PolyPhen-2^8^ scores were obtained through the web portal at http://genetics.bwh.harvard.edu/pph2/bgi.shtml. Positions of variants on the GPCR fold were determined using multiple alignments from GPCRdb^7^.

### Phenotype Definitions

#### Hospital Record and Verbal Questionnaire

As previously described, we used the following procedure to define cases and controls for non-cancer phenotypes^65^. For a given phenotype, ICD-10 codes (Data-Field 41202) were grouped with self-reported non-cancer illness codes from verbal questionnaires (Data-Field 20002) that were closely related. This was done by first creating a computationally generated candidate list of closely related ICD-10 codes and self-reported non-cancer illness codes, then manually curating the matches. The computational mapping was performed by calculating the token set ratio between the ICD-10 code description and the self-reported illness code description using the FuzzyWuzzy python package. The high-scoring ICD-10 matches for each self-reported illness were then manually curated to ensure high confidence mappings. Manual curation was required to validate the matches because fuzzy string matching may return words that are similar in spelling but not in meaning. For example, to create a hypertension cohort the code description from Data-Field 20002 (“Hypertension”) was mapped to all ICD-10 code descriptions and all closely related codes were returned (“I10: Essential (primary) hypertension” and “I95: Hypotension”). After manual curation code I10 would be kept and code I95 would be discarded. After matching ICD-10 codes and with self-reported illness codes, cases were identified for each phenotype using only the associated ICD-10 codes, only the associated self-reported illness codes, or both the associated ICD-10 codes and self-reported illness codes.

#### Family History

We used data from Category 100034 (Family history - Touchscreen - UK Biobank Assessment Centre) to define “cases” and controls for family history phenotypes. This category contains data from the touchscreen questionnaire on questions related to family size, sibling order, family medical history (of parents and siblings), and age of parents (age of death if died). We focused on Data Coding 20107: Illness of father and 20110: Illness of mother.

### Genetic Association Analyses

#### Quantitative phenotypes

We identified 146 quantitative phenotypes with at least 3,000 observations among the 337,205 white British subjects in the UK Biobank. As previously described, we took non-NA median of multiple measurements across up to three time points^66^. We focused on food intake, immune cell measurements, gross body measurements, behavioral phenotypes, and several other phenotypes (Table S1). We performed linear regression association analysis with v2.00a (20 Sep, 2017) for these 146 quantitative phenotypes. We performed quantile normalization for each phenotype (--pheno-quantile-normalize option), where we fit the linear model with covariates and transform the residuals to Normal distribution N(0, 1) while preserving the original rank in the residuals. We used the following covariates in our analysis: age, sex, array type, and the first four principal components, where array type is a binary variable that represents whether an individual was genotyped with UK Biobank Axiom Array or UK BiLEVE Axiom Array. For variants that were specific to one array, we did not use array as a covariate. We corrected p-values using the Benjamin-Yekutieli approach implemented in R’s p.adjust. We considered associations with BY-corrected p-values less than 0.05 as significant which controls the false discovery rate at 5%^67^.

#### Binary phenotypes

We identified 129 binary phenotypes with at least 2,000 cases among the 337,205 white British subjects in the UK Biobank and performed logistic regression association analysis with Firth-fallback using PLINK v2.00a (17 July 2017)^68^. Firth-fallback is a hybrid algorithm which normally uses the logistic regression code described in ^69^, but switches to a port of logistf() (https://cran.r-project.org/web/packages/logistf/index.html) in two cases: (1) one of the cells in the 2×2 allele count by case/control status contingency table is empty (2) logistic regression was attempted since all the contingency table cells were nonzero, but it failed to converge within the usual number of steps. We used the following covariates in our analysis: age, sex, array type, and the first four principal components, where array type is a binary variable that represents whether an individual was genotyped with UK Biobank Axiom Array or UK BiLEVE Axiom Array. For variants that were specific to one array, we did not use array as a covariate. We corrected p-values using the Benjamin-Yekutieli approach implemented in R’s p.adjust. We considered associations with BY-corrected p-values less than 0.05 as significant which controls the false discovery rate at 5%.

#### Comparison to GWAS Catalog

We compared significant associations to the GWAS Catalog^16^ to determine whether the associations we identified had been previously reported. We downloaded the GWAS Catalog v1.0.2 on August 23, 2019. We matched phenotypes in the GWAS Catalog to our phenotypes using the FuzzyWuzzy python package and manually reviewed matches. We erred on the side of including similar phenotypes from the GWAS Catalog for matches that were not perfect. We then matched the associated variants to variants in the GWAS Catalog using rs identifiers and chromosome:position identifiers. In order to match variants using chromosome:position, we used dbSNP or UCSC LiftOver^70^ to convert UK Biobank genotyping array coordinates in hg19 to GRCh38 coordinates to match with the GWAS Catalog.

We used LDlink (https://ldlink.nci.nih.gov/)^62^ to identify variants in LD with the variants that we found associations for using the GBR British population from 1000 Genomes Phase 3 (Version 5). We matched these variants in LD with the associated variants to variants in the GWAS Catalog using rs identifiers and chromosome:position identifiers when rs identifiers failed.

#### *ADRB2* Conditional Analyses

We fit a model for each of eosinophil counts, eosinophil percentage, monocyte percentage, and neutrophil count with the genotypes of rs1042713, rs1042714, and rs1800888 as independent variables. We included a constant and age, sex, array type, and the first four principal components as covariates for these models.

#### Coffee and Tea Intake Additivity Analysis

We fit two models for each of coffee and tea intake to test whether the associations were consistent with non-additive effects. We performed quantile normalization for each phenotype to transform the values to standard normal distribution. For the additive model, we encoded genotypes as 0,1,2 and included a single term for the effect of genotype on tea/coffee intake. For the genotypic model we included a term to indicate whether a subject was heterozygous and a separate term to indicate whether a subject was homozygous alternate. We included a constant and age, sex, array type, and the first four principal components as covariates for these regressions.

### *GIPR* characterization

#### Constructions

pcDNA3.1(+)_FLAG-hGIPR(WT) was produced by ligation of a Flag-tagged codon optimized GIPR insert in a pcDNA3.1(+) vector. The FLAG-GIPR insert was generated by PCR using the forward primer AGGATGACGACGATAAGGGGTACCACCAGCCCCATCCTGCAGC and the reverse primer TGGCAACTAGAAGGCACAGTC on a codon optimized ORF of GIPR (isoform 1, which has 466 amino acids) kindly given by Domain Therapeutics NA Inc (Canada, Montreal). For the pcDNA3.1(+) backbone, we digested pcDNA3.1(+)_FLAG-hMC4R(WT), kindly given by the laboratory of Pr Sadaaf Farooqi^71^, with KpnI-HF and XbaI. Fragments were then ligated using pEASY-Uni kit (CU101, Transgen Biotech) Then, pcDNA3.1(+)_FLAG-hGIPR(E354Q) was generated using overlap extension PCR protocol on the pcDNA3.1(+)_FLAG-hGIPR(WT) vector using the two primers described above in combination with respectively mutagenesis reverse primer ACACCACCTGGTGCACGCCCAGCAG and mutagenesis forward primer CGTGCACCAGGTGGTGTTTGCTCC.

#### BRET

Morphologically selected human embryonic kidney HEK293SL cells (Beautrait, Paradis et al. 2017) were cultured in Dulbecco’s modified Eagle medium (DMEM, Wisent) with 10% newborn calf serum iron supplemented (Wisent) and 1% penicillin/streptomycin (Wisent) at 37°C with 5% CO_2_. About 48h hours before experiments, 35 000 cells were seeded per well in white 96-well CellStar© plates (Greiner Bio-One) with transfection mix composed of 100 ng of 25 kDa linear polyethylenimine (Polysciences) and DNA for GRK2-based Gα activation biosensor (40 ng of FLAG-hGIPR(WT) or 45 ng of FLAG-hGIPR(E354Q), 60 ng of acceptor GRK2-GFP10, 1 ng of donor RlucII-Gγ5, 10 ng of Gβ1, and 10 ng of the designated Gα, completed to 126 ng total with empty pcDNA3.1(+) vector) or βarr2 biosensor (40 ng of FLAG-hGIPR, 40 ng of acceptor rGFP-CAAX, 6 ng of donor βarr2-RlucII and 14 ng of empty pcDNA3.1(+) vector). The day of the experiment, cells were washed with Dulbecco’s phosphate buffered saline (D-PBS, Wisent) supplemented with 1 mM CaCl_2_ and 1 mM MgCl_2_ (CM), and then put in Tyrode HEPES buffer (THB: 137 mM NaCl, 25 mM HEPES, 11,9 mM NaHCO_3_, 5,5 mM D-glucose, 3,6 mM NaH_2_PO_4_, 1 mM CaCl_2_, 1 mM MgCl_2_ and 0,9 mM KCl; pH adjusted to 7,4 and filtered at 0,22 micron). After incubation of at least 1 hour to stabilize the signal, cells were stimulated with increasing concentrations of agonist GIP (RP10795, GenScript) for 45 minutes. Five minutes before reading, luciferase substrate’s coelenterazine 400a DeepBlueC™ (Nanoligth technology) was added to the final concentration of 2,5 µM. Luminescence signal was red on TriStar^2^ multidetection microplate reader (LB942, Berthold Technologies) at 410/80 nm and 515/40 nm and integrated for 0,1 second. Plates were kept and red at 37°C. Dose-response curves were fitted in GraphPad Prism 8 using three parameters non-linear logistic regression to estimate top and bottom asymptotes. Results are expressed as fold over the bottom asymptote of the WT receptor. Maximal stimulation for WT receptor was set to 100 and E_max_ for E354Q variant was calculated as % of the range of the WT response E_max_ EQ = (top asymptote EQ – bottom asymptote WT) / span WT * 100. Statistical analysis was done using T-test function of Excel for heteroscedastic data using bilateral distribution (unequal variance confirmed with F-test) on EC_50_ and E_max_ of each individual experiment (N=2 for Gz; N=3 for Gs, Gi2, GoB and G15; and N=4 for β-arrestin2). In text data are expressed as mean ± SD.

#### Cell-surface ELISA

The same pools of cells used for BRET were seeded on transparent 96-well Costar© plates (Corning), and cells transfected with empty vector pcDNA3.1(+) were used as no receptor condition. The day of experiment, cells were washed with D-PBS+CM and fixed with 50 µL of 3% para-formaldehyde (PFA) in D-PBS+CM for then minutes. PFA was discarded and cells were stored in D-PBS+CM at 4°C until the rest of the experiment was carried. Cells were washed with 3 × 100 µL of 2% bovine serum albumin (BSA, Sigma) in D-PBS+CM and blocked in that same solution for 2 hours at room temperature (RT). Monoclonal ANTI-FLAG® M2-Peroxidase (HRP) antibody (Sigma) was then added at dilution of 1/1000 in 2% BSA for 2 hours at RT. Cells were washed again with 3 × 100 µL of 2% BSA, then with 3 × 100 µL of D-PBS+CM. Then, cells were incubated with 100 µL of a solution of SigmaFast™ OPD (Sigma) for 30 minutes at RT. For Gα activation experiments, 100 µL of the supernatant were transferred in another transparent plate before reading absorbance at 450 ± 2 nm on SpectraMax® 190 microplate reader (Molecular Devices). For β-arrestin2 recruitment assays, 25 µL of HCl 3N were added to stop the reaction, and absorbance was red at 492 ± 2 nm. Cell-surface expression as fold of the WT was calculated as the ratio of the specific counts of the E354Q mutant over those of the WT receptor = (OPD _EQ_ – OPD _no receptor_) / (OPD _WT_ – OPD _no receptor_).

## Supplementary Information

### Supplementary Figures

**Figure S1.**
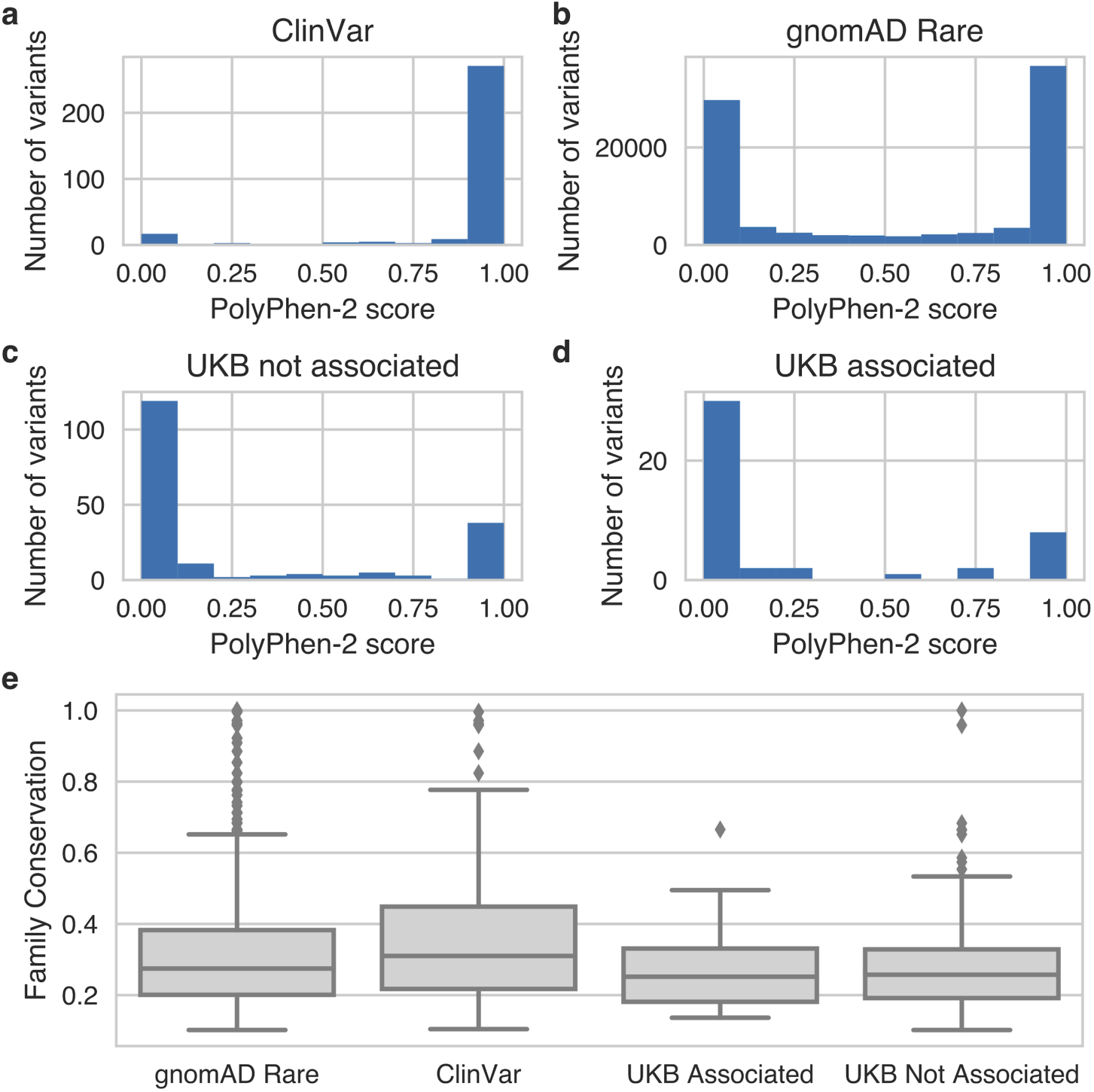
Distribution of PolyPhen-2 scores for (a) GPCR variants in ClinVar; (b) rare (MAF<1%) GPCR variants in gnomAD, (c) variants analyzed in this study that were not significantly associated with any phenotype, and (d) variants analyzed in this study that were significantly associated with at least one phenotype. (e) Distribution of family conservation scores for variants in class A GPCRs across the four groups of variants in a-d. The distribution of conservation scores is significantly different (Wilcoxon p<0.05) between the gnomAD Rare and ClinVar variants and between the ClinVar and UKB Not Associated variants.

**Figure S2.**
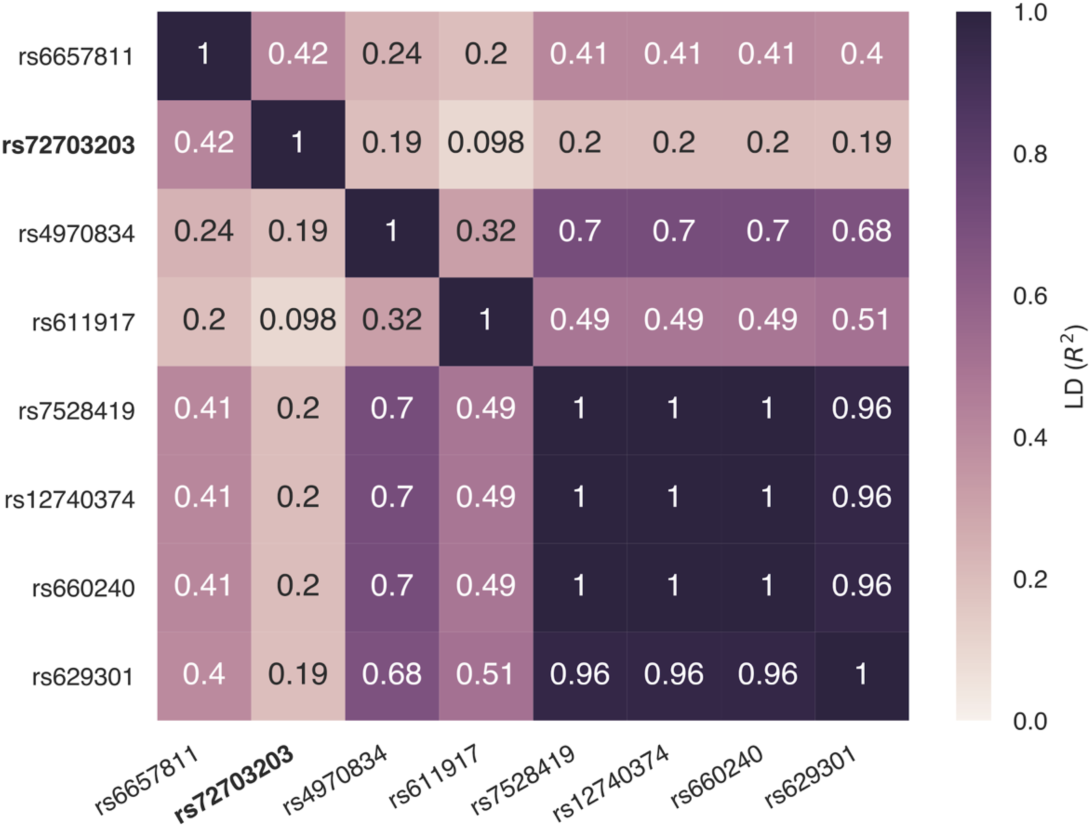
Linkage disequilibrium in the GBR British population between rs72703203 (bold) and other *CELSR2* variants reported in the GWAS Catalog as significantly associated with various lipid and cardiac phenotypes.

**Figure S3.**
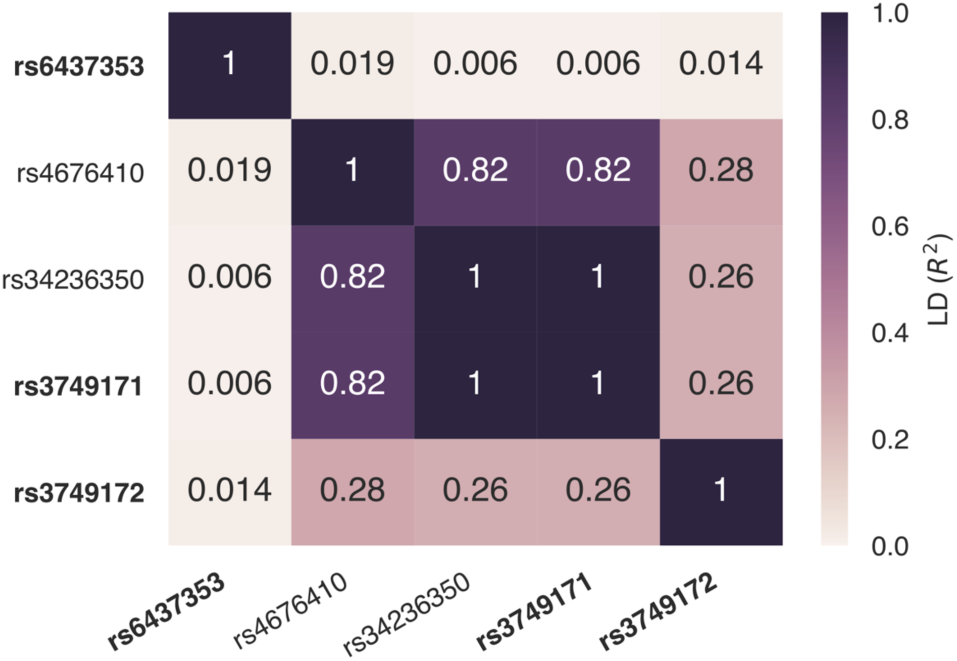
Linkage disequilibrium in the GBR British population between variants in *GPR35* with significant associations in this study (bold) and variants reported in the GWAS Catalog.

**Figure S4.**
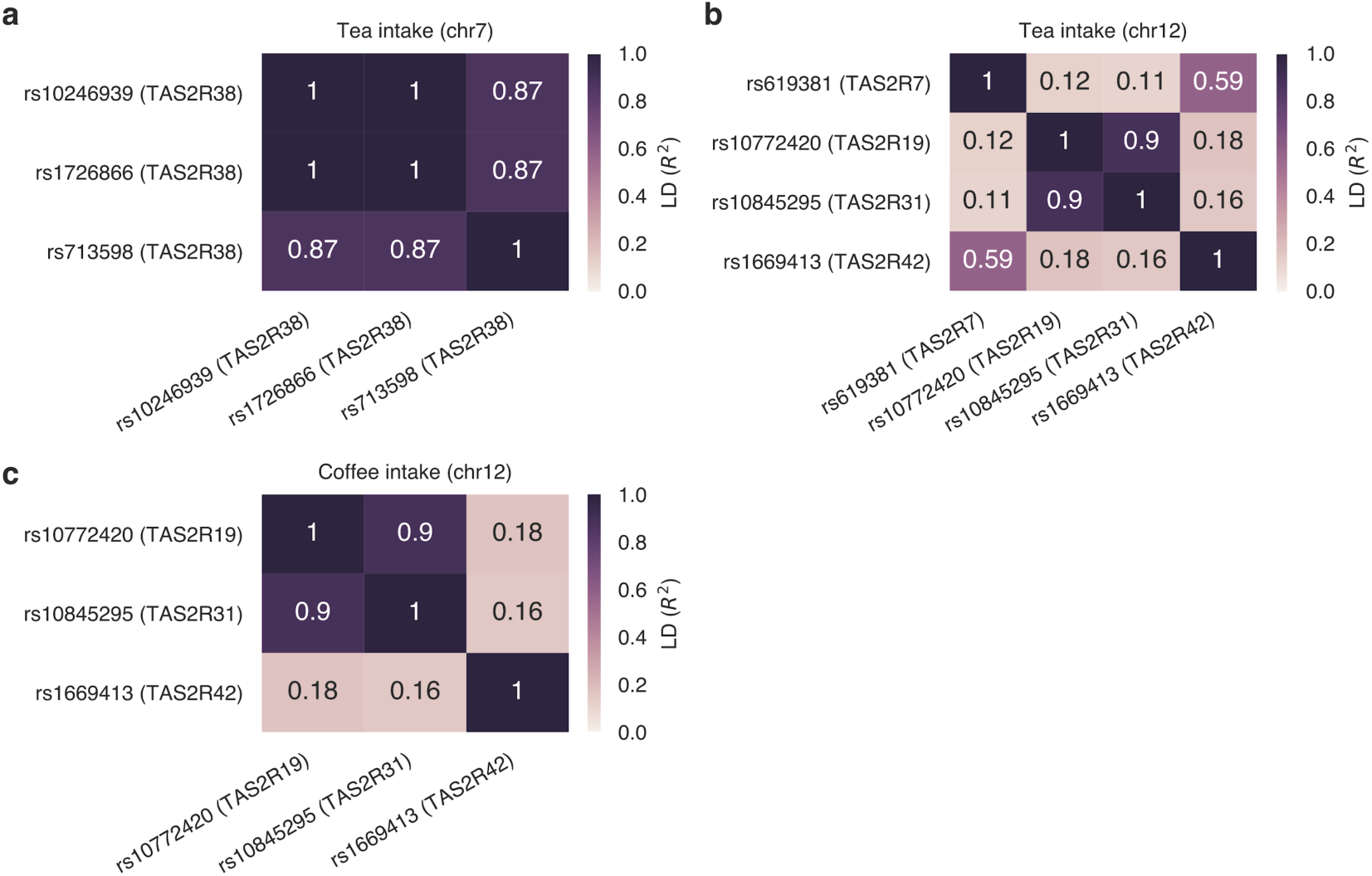
Linkage disequilibrium in the GBR British population between variants in taste receptors associated with (a) tea intake on chromosome 7, (b) tea intake on chromosome 12, or (c) coffee intake on chromosome 12.

**Figure S5.**
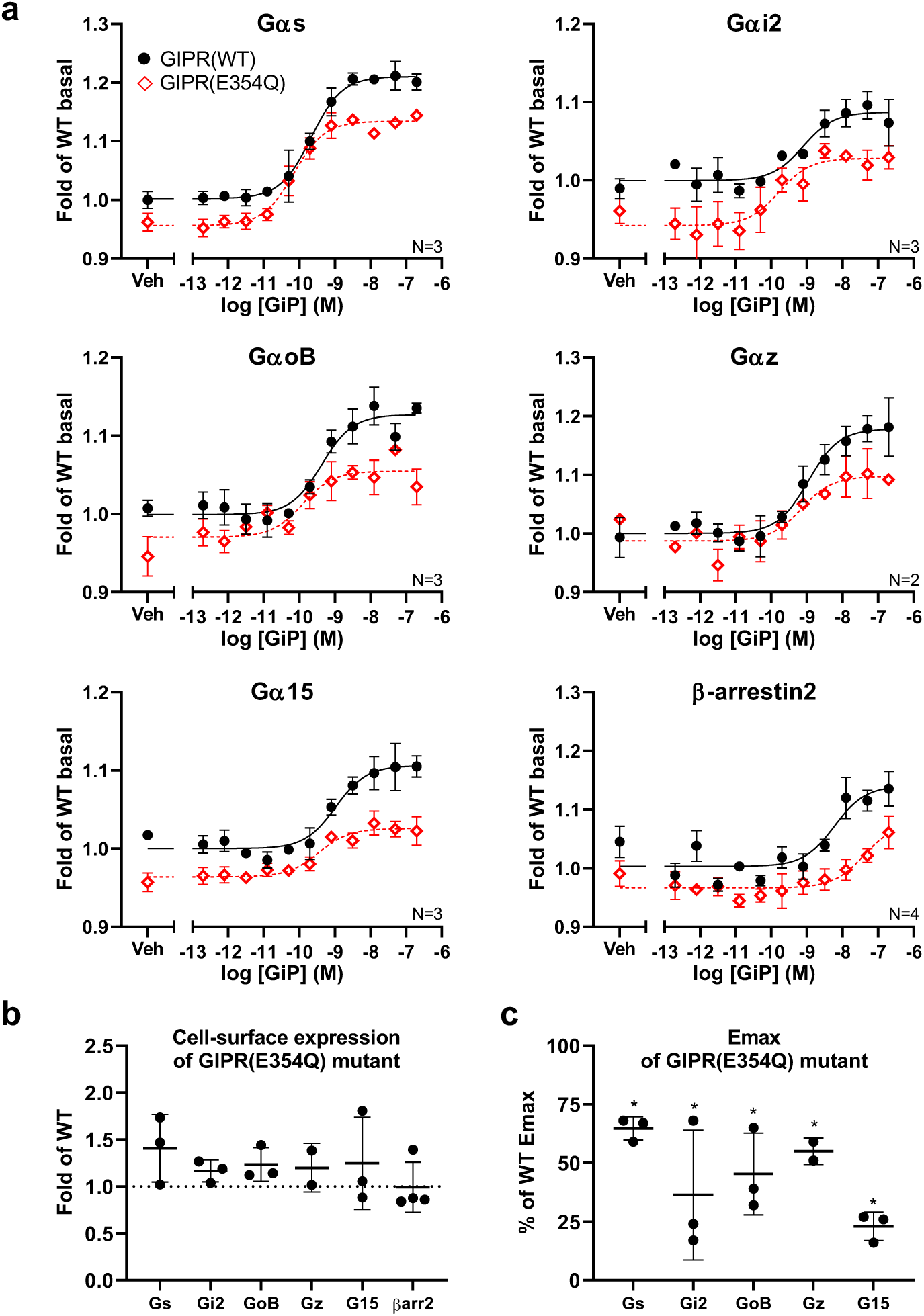
Characterization of the signaling profile of *GIPR* rs1800437 (p.Glu354Gln, E354Q) reference (Glu) and alternate (Gln) alleles. (a) Dose-response curves for GIP stimulation using BRET-based assay for Gα activation or β-arrestin2 recruitment. Data are expressed as mean ± SEM if the number of independent experiments (N) is ≥3, or mean ± SD if N<3. EC50 were not statistically different. Corresponding (b) cell-surface expression and (c) Emax of the p.Glu354Gln alternate allele (E354Q mutant) for the experiments shown in (a). Each dot represents the value for an independent experiment (the mean of at least three replicates for (b), or the calculated Emax value for (c)), and the error bars represent the mean ± standard deviation of all independent experiments. Emax values for β-arrestin2 are not shown since saturation was not always reached. * = *p*-value < 0.05, t-test on EC50 and Emax of each individual experiment.

**Figure S6.**
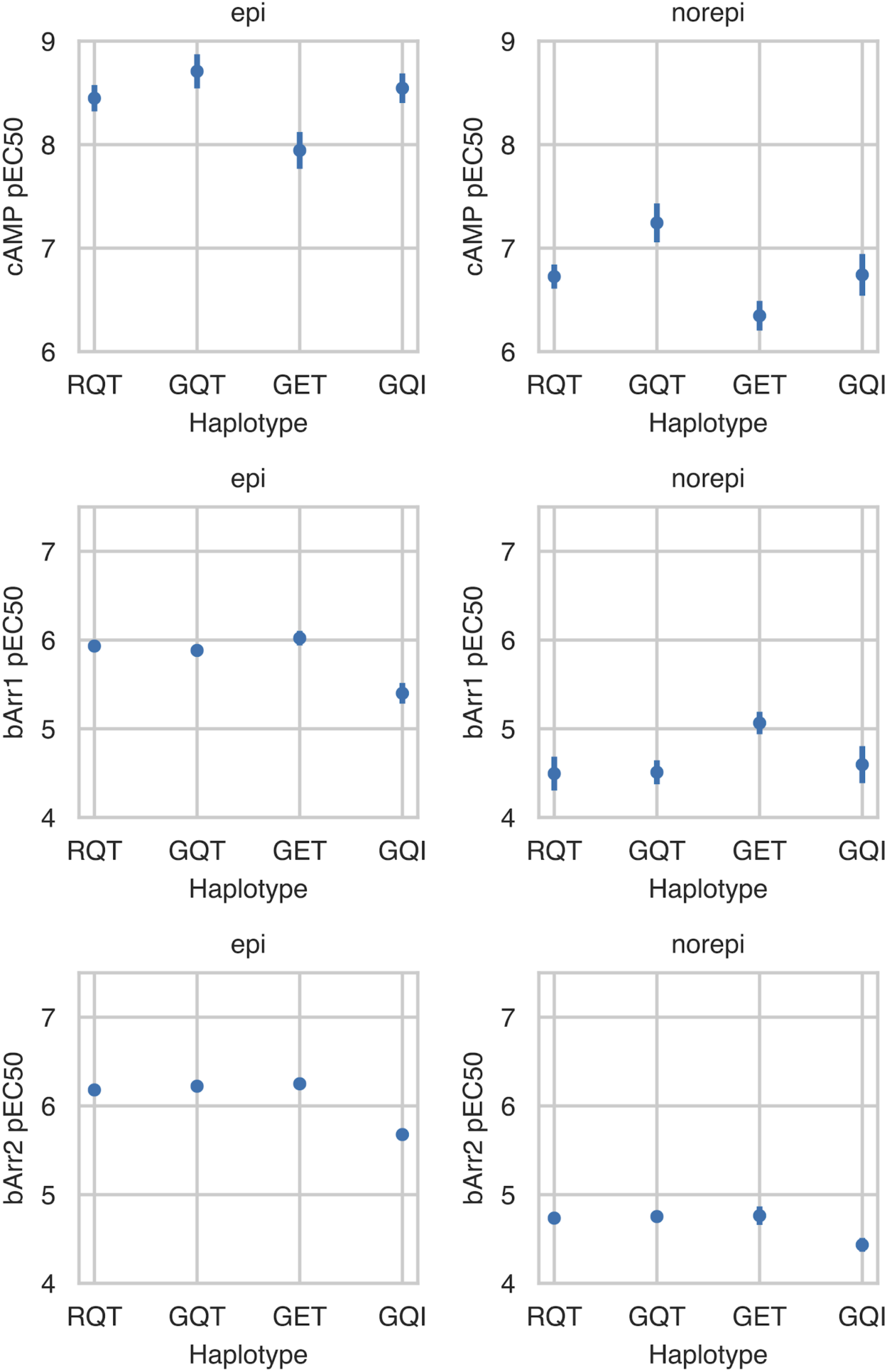
pEC50 for cAMP signaling, beta-arrestin-1 recruitment, and beta-arrestin-2 recruitment for four *ADRB2* haplotypes upon stimulation with the endogenous ligands epinephrine (epi) and norepinephrine (norepi). Tukey’s test p-values for all pairwise comparisons are reported in Table S4.

### Supplementary Tables

Table S1. Phenotypes and GPCR genes and variants used for PheWAS. The “GPCR genes” tab contains 398 GPCR genes that we used to identify GPCR genetic variants. The “GPCR tested variants” tab contains the 268 variants that we tested in the PheWAS analysis. The “GPCR variant summary” tab shows the number of each variant type included in the pheWAS analysis. The “Quantitative phenotypes” and “Binary phenotypes” tabs contain the phenotypes analyzed and the number of observations and cases, respectively.

Table S2. GPCR PheWAS results. Summary statistics for quantitative and binary associations with BY-adjusted p-value less than 0.05.

Table S3. GPCR variant annotations. Across-family conservation scores, GPCR functional region annotations, and PolyPhen-2 pathogenicity scores for GPCR variants assessed in this study, 86,601 rare GPCR variants from gnomAD (MAF<1%) and 318 GPCR variants reported in ClinVar.

Table S4. p-values from Tukey’s test for differences in pEC50 for cAMP signaling, beta-arrestin-1 recruitment, and beta-arrestin-2 recruitment for four *ADRB2* haplotypes upon stimulation with the endogenous ligands epinephrine (epi) and norepinephrine (norepi).

## Acknowledgments

This research has been conducted using the UK Biobank Resource under Application Number 24983, “Generating effective therapeutic hypotheses from genomic and hospital linkage data” (http://www.ukbiobank.ac.uk/wp-content/uploads/2017/06/24983-Dr-Manuel-Rivas.pdf). Based on the information provided in Protocol 44532 the Stanford IRB has determined that the research does not involve human subjects as defined in 45 CFR 46.102(f) or 21 CFR 50.3(g). All participants of UK Biobank provided written informed consent (more information is available at https://www.ukbiobank.ac.uk/2018/02/gdpr/). This work was supported by National Human Genome Research Institute (NHGRI) of the National Institutes of Health (NIH) under award R01HG010140 (M.A.R.), and by NIH grant R01 GM127359 (to R.O.D.). The content is solely the responsibility of the authors and does not necessarily represent the official views of the National Institutes of Health. M.A.R. is supported by Stanford University and a National Institute of Health center for Multi- and Trans-ethnic Mapping of Mendelian and Complex Diseases grant (5U01 HG009080). C.D. and A.J.V are supported by the Stanford ChEM-H Postdocs at the Interface Seed Grant. J.M.P. is supported by a Stanford Graduate Fellowship. S.A.L. is supported by the Ph.D scholarship of the Institute for Research in Cancer and Immunology. The authors would like to thank the Genome Aggregation Database (gnomAD) and the groups that provided exome and genome variant data to this resource. A full list of contributing groups can be found at https://gnomad.broadinstitute.org/about.

## Author Contributions

C.D., A.J.V., R.O.D., and M.A.R. conceived and designed the study. C.D., A.J.V., J.M.P., Y.T., G.V., and M.A.R. designed and carried out the statistical and computational analyses. F.M.H., S.A.L., and M.B. designed and carried out *GIPR* and *ADRB2* experiments. F.M.H., S.A.L., M.M., M.B., A.J.V, and R.O.D. interpreted the results. The manuscript was written by C.D., A.J.V., J.M.P., S.A.L., M.B., R.O.D., and M.A.R. R.O.D. and M.A.R. supervised all aspects of the study.

## References

1. Katritch, V., Cherezov, V. & Stevens, R. C. Structure-function of the G protein-coupled receptor superfamily. Annu. Rev. Pharmacol. Toxicol. 53, 531–556 (2013).

2. Sudlow, C. et al. UK biobank: an open access resource for identifying the causes of a wide range of complex diseases of middle and old age. PLoS Med. 12, e1001779 (2015).

3. Altshuler, D., Daly, M. J. & Lander, E. S. Genetic mapping in human disease. Science 322, 881–888 (2008).

4. Venkatakrishnan, A. J. et al. Molecular signatures of G-protein-coupled receptors. Nature vol. 494 185–194 (2013).

5. Dong, X., Han, S., Zylka, M. J., Simon, M. I. & Anderson, D. J. A diverse family of GPCRs expressed in specific subsets of nociceptive sensory neurons. Cell 106, 619–632 (2001).

6. Divorty, N., Mackenzie, A. E., Nicklin, S. A. & Milligan, G. G protein-coupled receptor 35: an emerging target in inflammatory and cardiovascular disease. Front. Pharmacol. 6, 19189 (2015).

7. Pándy-Szekeres, G. et al. GPCRdb in 2018: adding GPCR structure models and ligands. Nucleic Acids Res. 46, D440–D446 (2018).

8. Adzhubei, I. A. et al. A method and server for predicting damaging missense mutations. Nat. Methods 7, 248–249 (2010).

9. Genome Aggregation Database Consortium et al. The mutational constraint spectrum quantified from variation in 141,456 humans. Nature 581, 434–443 (2020).

10. Landrum, M. J. et al. ClinVar: improving access to variant interpretations and supporting evidence. Nucleic Acids Res. 46, D1062–D1067 (2018).

11. Rosen, M. D. & Privalsky, M. L. Thyroid hormone receptor mutations in cancer and resistance to thyroid hormone: perspective and prognosis. J. Thyroid Res. 2011, 361304 (2011).

12. Pinato, D. J. & Mauri, F. A. Metastatic Spread of Lung Cancer to Brain and Liver. in Brain Metastases from Primary Tumors 123–129 (2014).

13. Astle, W. J. et al. The Allelic Landscape of Human Blood Cell Trait Variation and Links to Common Complex Disease. Cell 167, 1415–1429.e19 (2016).

14. Chan, I. H. & Privalsky, M. L. Thyroid hormone receptor mutants implicated in human hepatocellular carcinoma display an altered target gene repertoire. Oncogene 28, 4162–4174 (2009).

15. Lotta, L. A. et al. Human Gain-of-Function MC4R Variants Show Signaling Bias and Protect against Obesity. Cell 177, 597–607.e9 (2019).

16. Buniello, A. et al. The NHGRI-EBI GWAS Catalog of published genome-wide association studies, targeted arrays and summary statistics 2019. Nucleic Acids Research vol. 47 D1005–D1012 (2019).

17. International Headache Genetics Consortium et al. Meta-analysis of 375,000 individuals identifies 38 susceptibility loci for migraine. Nat. Genet. 48, 856–866 (2016).

18. Meng, W. et al. A Genome-Wide Association Study Finds Genetic Associations with Broadly-Defined Headache in UK Biobank (N = 223,773). EBioMedicine 28, 180–186 (2018).

19. Manteniotis, S. et al. Comprehensive RNA-Seq expression analysis of sensory ganglia with a focus on ion channels and GPCRs in Trigeminal ganglia. PLoS One 8, e79523 (2013).

20. Zhang, L. et al. Cloning and expression of MRG receptors in macaque, mouse, and human. Brain Res. Mol. Brain Res. 133, 187–197 (2005).

21. Gourraud, P.-A. et al. A genome-wide association study of brain lesion distribution in multiple sclerosis. Brain 136, 1012–1024 (2013).

22. Raffield, L. M. et al. Heritability and genetic association analysis of neuroimaging measures in the Diabetes Heart Study. Neurobiol. Aging 36, 1602.e7–15 (2015).

23. Pujol-Borrell, R., Giménez-Barcons, M., Marín-Sánchez, A. & Colobran, R. Genetics of Graves’ Disease: Special Focus on the Role of TSHR Gene. Horm. Metab. Res. 47, 753–766 (2015).

24. van der Harst, P. & Verweij, N. Identification of 64 Novel Genetic Loci Provides an Expanded View on the Genetic Architecture of Coronary Artery Disease. Circ. Res. 122, 433–443 (2018).

25. Postmus, I. et al. Pharmacogenetic meta-analysis of genome-wide association studies of LDL cholesterol response to statins. Nat. Commun. 5, 5068 (2014).

26. Arvind, P., Nair, J., Jambunathan, S., Kakkar, V. V. & Shanker, J. CELSR2–PSRC1– SORT1 gene expression and association with coronary artery disease and plasma lipid levels in an Asian Indian cohort. J. Cardiol. 64, 339–346 (2014).

27. Grallert, H. et al. Eight genetic loci associated with variation in lipoprotein-associated phospholipase A2 mass and activity and coronary heart disease: meta-analysis of genome-wide association studies from five community-based studies. Eur. Heart J. 33, 238–251 (2012).

28. Ma, L. et al. Genome-wide association analysis of total cholesterol and high-density lipoprotein cholesterol levels using the Framingham heart study data. BMC Med. Genet. 11, 55 (2010).

29. Musunuru, K. & Kathiresan, S. Genetics of Common, Complex Coronary Artery Disease. Cell 177, 132–145 (2019).

30. Takata, R. et al. Genome-wide association study identifies five new susceptibility loci for prostate cancer in the Japanese population. Nat. Genet. 42, 751–754 (2010).

31. Hoffmann, T. J. et al. A Large Multiethnic Genome-Wide Association Study of Prostate Cancer Identifies Novel Risk Variants and Substantial Ethnic Differences. Cancer Discovery vol. 5 878–891 (2015).

32. Wang, M. et al. Large-scale association analysis in Asians identifies new susceptibility loci for prostate cancer. Nat. Commun. 6, 8469 (2015).

33. Tang, X.-L., Wang, Y., Li, D.-L., Luo, J. & Liu, M.-Y. Orphan G protein-coupled receptors (GPCRs): biological functions and potential drug targets. Acta Pharmacologica Sinica vol. 33 363–371 (2012).

34. (igas), I. G. of A. S. C. & International Genetics of Ankylosing Spondylitis Consortium (IGAS). Identification of multiple risk variants for ankylosing spondylitis through high-density genotyping of immune-related loci. Nature Genetics vol. 45 730–738 (2013).

35. Venkateswaran, S. et al. Enhanced Contribution of HLA in Pediatric Onset Ulcerative Colitis. Inflamm. Bowel Dis. 24, 829–838 (2018).

36. The International IBD Genetics Consortium (IIBDGC) et al. Analysis of five chronic inflammatory diseases identifies 27 new associations and highlights disease-specific patterns at shared loci. Nat. Genet. 48, 510–518 (2016).

37. Ellinghaus, D. et al. Genome-wide association analysis in Primary sclerosing cholangitis and ulcerative colitis identifies risk loci at GPR35 and TCF4. Hepatology 58, 1074–1083 (2013).

38. International Multiple Sclerosis Genetics Consortium et al. Association analyses identify 38 susceptibility loci for inflammatory bowel disease and highlight shared genetic risk across populations. Nat. Genet. 47, 979–986 (2015).

39. The International IBD Genetics Consortium (IIBDGC) et al. Host–microbe interactions have shaped the genetic architecture of inflammatory bowel disease. Nature 491, 119–124 (2012).

40. Li, Y. R. et al. Meta-analysis of shared genetic architecture across ten pediatric autoimmune diseases. Nat. Med. 21, 1018–1027 (2015).

41. CHARGE-Heart Failure Consortium et al. Trans-ancestry meta-analyses identify rare and common variants associated with blood pressure and hypertension. Nat. Genet. 48, 1151–1161 (2016).

42. Hayes, J. E. et al. Allelic Variation in TAS2R Bitter Receptor Genes Associates with Variation in Sensations from and Ingestive Behaviors toward Common Bitter Beverages in Adults. Chem. Senses 36, 311–319 (2011).

43. Hayes, J. E., Feeney, E. L., Nolden, A. A. & McGeary, J. E. Quinine Bitterness and Grapefruit Liking Associate with Allelic Variants in TAS2R31. Chem. Senses 40, 437–443 (2015).

44. Kim, U.-K. Positional Cloning of the Human Quantitative Trait Locus Underlying Taste Sensitivity to Phenylthiocarbamide. Science 299, 1221–1225 (2003).

45. Risso, D. S. et al. Global diversity in the TAS2R38 bitter taste receptor: revisiting a classic evolutionary PROPosal. Sci. Rep. 6, 115 (2016).

46. Dupre, J., Ross, S. A., Watson, D. & Brown, J. C. STIMULATION OF INSULIN SECRETION BY GASTRIC INHIBITORY POLYPEPTIDE IN MAN. 1. J. Clin. Endocrinol. Metab. 37, 826–828 (1973).

47. Miyawaki, K. et al. Glucose intolerance caused by a defect in the entero-insular axis: A study in gastric inhibitory polypeptide receptor knockout mice. Proceedings of the National Academy of Sciences vol. 96 14843–14847 (1999).

48. Namkung, Y. et al. Monitoring G protein-coupled receptor and β-arrestin trafficking in live cells using enhanced bystander BRET. Nature Communications vol. 7 (2016).

49. Ehrlich, A. T. et al. Biased Signaling of the Mu Opioid Receptor Revealed in Native Neurons. iScience vol. 14 47–57 (2019).

50. Avet, C. et al. Selectivity Landscape of 100 Therapeutically Relevant GPCR Profiled by an Effector Translocation-Based BRET Platform. Pharmacology and Toxicology D1006 (2020).

51. andMe Research Team et al. Gene discovery and polygenic prediction from a genome-wide association study of educational attainment in 1.1 million individuals. Nat. Genet. 50, 1112–1121 (2018).

52. Kichaev, G. et al. Leveraging polygenic functional enrichment to improve GWAS power. doi: 10.1101/222265.

53. Thanawala, V. J. et al. β 2 -Adrenoceptor Agonists Are Required for Development of the Asthma Phenotype in a Murine Model. Am. J. Respir. Cell Mol. Biol. 48, 220–229 (2013).

54. Moore, C. A. C., Milano, S. K. & Benovic, J. L. Regulation of Receptor Trafficking by GRKs and Arrestins. Annu. Rev. Physiol. 69, 451–482 (2007).

55. Fortin, J.-P., Schroeder, J. C., Zhu, Y., Beinborn, M. & Kopin, A. S. Pharmacological Characterization of Human Incretin Receptor Missense Variants. J. Pharmacol. Exp. Ther. 332, 274–280 (2010).

56. Kubota, A. et al. Identification of two missense mutations in the GIP receptor gene: a functional study and association analysis with NIDDM: no evidence of association with Japanese NIDDM subjects. Diabetes vol. 45 1701–1705 (1996).

57. Mohammad, S. et al. A Naturally Occurring GIP Receptor Variant Undergoes Enhanced Agonist-Induced Desensitization, Which Impairs GIP Control of Adipose Insulin Sensitivity. Mol. Cell. Biol. 34, 3618–3629 (2014).

58. Almind, K. et al. Discovery of amino acid variants in the human glucose-dependent insulinotropic polypeptide (GIP) receptor: the impact on the pancreatic beta cell responses and functional expression studies in Chinese hamster fibroblast cells. Diabetologia vol. 41 1194–1198 (1998).

59. Gabe, M. B. N. et al. Enhanced agonist residence time, internalization rate and signalling of the GIP receptor variant [E354Q] facilitate receptor desensitization and long-term impairment of the GIP system. Basic Clin. Pharmacol. Toxicol. 70, 14.1- (2019).

60. Costa-Neto, C. M., Parreiras-e-Silva, L. T. & Bouvier, M. A Pluridimensional View of Biased Agonism. Molecular Pharmacology vol. 90 587–595 (2016).

61. McLaren, W. et al. The Ensembl Variant Effect Predictor. Genome Biol. 17, S8 (2016).

62. Machiela, M. J. & Chanock, S. J. LDlink: a web-based application for exploring population-specific haplotype structure and linking correlated alleles of possible functional variants: Fig. 1. Bioinformatics vol. 31 3555–3557 (2015).

63. Church, D. M. et al. Modernizing Reference Genome Assemblies. (2011) doi: 10.1371/journal.pbio.1001091.

64. Karczewski, K. J. et al. Variation across 141,456 human exomes and genomes reveals the spectrum of loss-of-function intolerance across human protein-coding genes. Genomics 806 (2019).

65. DeBoever, C. et al. Medical relevance of protein-truncating variants across 337,205 individuals in the UK Biobank study. Nat. Commun. 9, 1066 (2018).

66. Tanigawa, Y. et al. Components of genetic associations across 2,138 phenotypes in the UK Biobank highlight adipocyte biology. Nat. Commun. 10, 1001 (2019).

67. Yekutieli, D. & Benjamini, Y. under dependency. Ann. Stat. 29, 1165–1188 (2001).

68. Chang, C. C. et al. Second-generation PLINK: rising to the challenge of larger and richer datasets. GigaScience vol. 4 (2015).

69. Hill, A. et al. Stepwise Distributed Open Innovation Contests for Software Development: Acceleration of Genome-Wide Association Analysis. Gigascience 6, 1–10 (2017).

70. Kent, W. J. et al. The Human Genome Browser at UCSC. Genome Res. 12, 996–1006 (2002).

71. Farooqi, I. S. et al. Dominant and recessive inheritance of morbid obesity associated with melanocortin 4 receptor deficiency. Journal of Clinical Investigation vol. 106 271–279 (2000).

